# Generating the head direction signal: Two types of head direction cells in the lateral mammillary nuclei and dorsal tegmental nuclei

**DOI:** 10.1101/2025.04.11.648409

**Authors:** Jeffrey S. Taube, William N. Butler, Julie R. Dumont, Jalina A. Graham, Jennifer L. Marcroft, Michael E. Shinder, Robert W. Stackman, Ryan M. Yoder

**Author notes:** <u>Correspondence</u>: Jeffrey S. Taube, Dartmouth College, Department of Psychological & Brain Sciences, 6207 Moore Hall, Hanover, NH, USA 03755.

## Abstract

Head direction (HD) cells discharge as a function of the animal’s directional heading and are believed to underlie one’s sense of direction. They have been identified in several brain areas, although the signal is thought to be generated across the connections between the lateral mammillary (LMN) and dorsal tegmental nuclei (DTN). Computational models have proposed that a ring-attractor network underlies the mechanisms that generate the signal. These models usually contain separate populations of neurons that encode HD and angular head velocity (AHV). Currently, both cell types have been identified in the LMN and DTN. However, HD attractor models also require cells, referred to as ‘rotation’ cells, which are sensitive to both parameters conjunctively (HD+AHV). Here we sought to identify such cells in the LMN-DTN network. We identified two distinct types of HD cells. The majority of LMN HD cells (∼64%) were AHV-independent, responding only to the animal’s directional heading. However, a second population (∼36%) was sensitive to both HD and AHV. Both symmetric and asymmetric AHV cell types were found. Similar results were found in the DTN, but with a higher percentage of conjunctive HD+AHV cells (60%). Notably, many HD+AHV conjunctive cells were also sensitive to the animal’s linear velocity (LV). In contrast, HD cells in the anterodorsal thalamus were rarely sensitive to AHV or LV. These findings demonstrate that the requisite rotation-type HD cell is present in brain areas responsible for generating the HD signal and supports the view that an attractor style network underlies its generation in mammals.

**Statements and Declaratio:** This manuscript does not represent the official view of the National Institute of Neurological Disorders and Stroke (USA) (NINDS), the National Institutes of Health (USA) (NIH), or any part of the US Federal Government. No official support or endorsement of this article by the NINDS or NIH is intended or should be inferred.

## Introduction

Accurate perception of your directional heading is essential for successful navigation. Numerous studies have identified populations of cells within various brain areas that discharge as a function of the animal’s directional heading, independent of its location and on-going behavior. These cells, referred to as head direction (HD) cells were originally identified in the postsubiculum of rats (Taube et al., 1990), but have since been identified in many limbic system and cortical areas, as well as across many different species – from insects (Seelig & Jayaraman, 2015), to fish (Petrucco et al., 2023), to non-human primates (Laurens et al., 2016). How the HD signal is generated has been a subject of investigation for several decades now. Studies using a variety of approaches have demonstrated that an intact vestibular labyrinth is critical for the generation of the HD signal (Stackman & Taube, 1997; Muir et al., 2009; Valerio & Taube, 2016) – particularly in the anterodorsal thalamus (ADN), where HD cells are abundant (Taube, 1995). To date, lesions of the ADN have led to the loss of direction-specific firing in all cortical, striatal, and other limbic areas where HD cells are present (Cullen & Taube, 2017). Further, lesions of subcortical structures (supragenual nucleus, DTN, LMN) that connect the vestibular nuclei, either directly or indirectly, to the ADN impair direction-specific firing in the ADN (Bassett et al., 2007; Biazoli et al., 2006; Clark et al., 2012) and suggest that there is an overall serial structure to processing the HD signal. It is generated in subcortical areas, transmitted to the ADN, which is used as a key node, and then conveyed broadly to many cortical areas (Sharp et al., 2001a; Taube, 2007)(Fig 1B).

**Figure 1.**
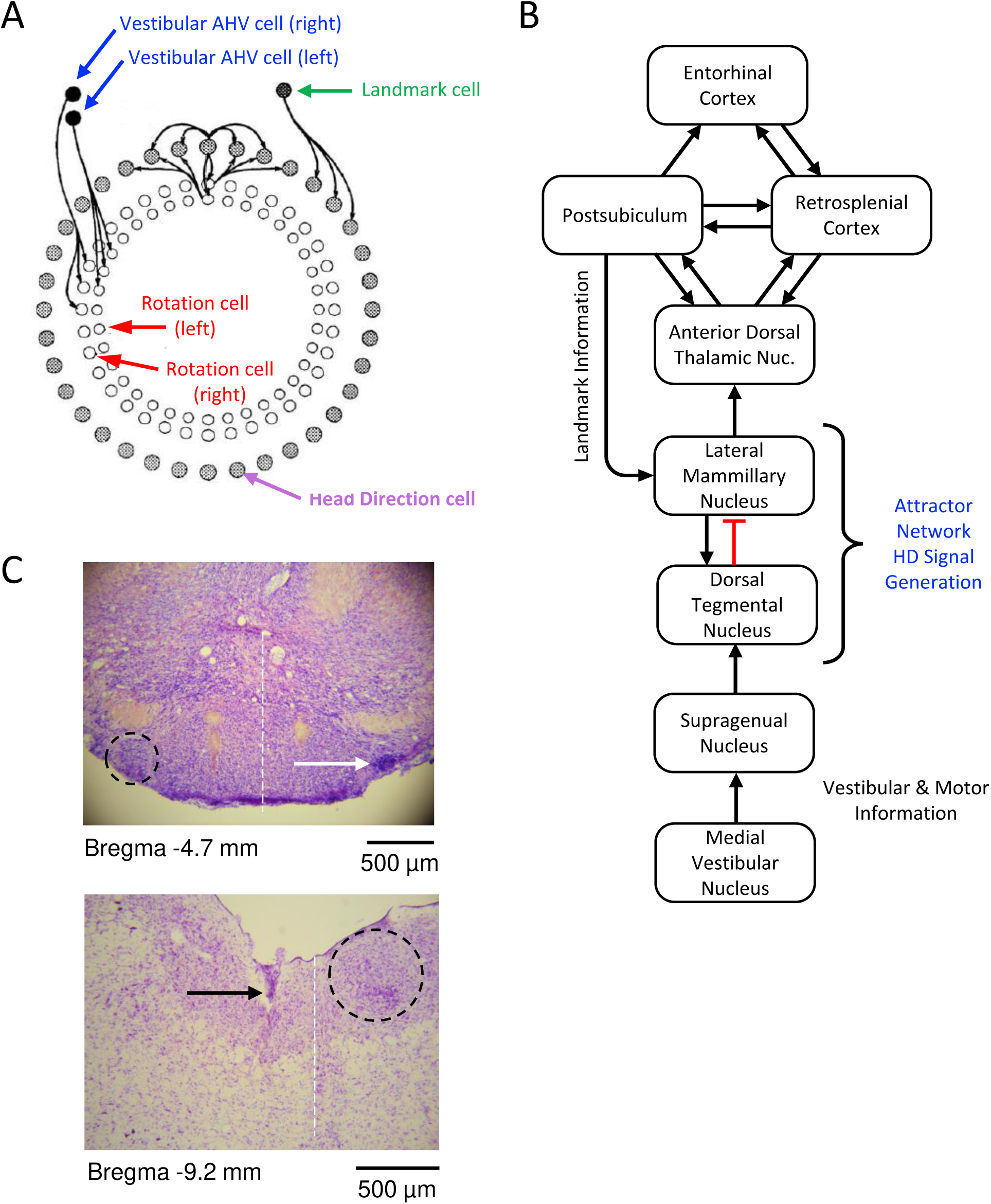
Generation of the HD signal. **A)** Ring attractor network model of how the head direction signal is generated. This model contains three cell rings. The head direction ring contains head direction cells representing different preferred firing directions and are not sensitive to AHV. HD cells project to nearby HD cells that represent adjacent HDs and are excitatory. They also project to rotation cells that represent similar HDs. Two additional rings contain rotation cells - one for CW turns and one for CCW turns. Each rotation cell is conjunctively sensitive to HD and AHV and receives inputs from both the HD cell ring and from vestibular inputs conveying AHV information. Rotation cells in turn project back to HD cells in the HD ring that are either CW or CCW shifted from the HD cell representing the current HD. Landmark cells, which connect directly to the HD ring convey spatial information about landmarks that are typically from visual sources, but can be from other types of sensory cues. Modified from Skaggs et al. 1995. **B)** Circuit diagram showing the major brain areas involved in generating the HD signal and conveying it to cortical areas. Vestibular and motor information originates from the vestibular nucleus and is projected rostrally to the supragenual nucleus and then to DTN and LMN. Landmark spatial information is processed in the cortex and is projected caudally from the postsubiculum to the LMN. All connections are excitatory except the pathway denoted in red. **C)** Histological images showing electrode tracks through the LMN (top) and DTN (bottom). Dashed circles encircle LMN and DTN on the side opposite where the electrode track can be seen (LMN: white arrow; DTN black arrow). For the LMN section, a Prussian blue reaction is evident where the electrode tips ended. The white dashed line indicates the midline in each section. The tissue to the left of the midline in the DTN section is a little deformed due to passage of the electrode array.

Studies that have investigated the generation of the HD signal have conceptually focused on models where a network of neurons is inter-connected in a ring-attractor structure (Skaggs et al., 1995; Redish et al., 1996; Zhang, 1996). The animal’s current HD is represented by a ‘hill of activity’ on the ring that moves to a new orientation when the animal makes a head turn. If an attractor type network underlies the formation of the HD signal, the next question is where this network resides in the brain. In rodents there are several lines of evidence that support the view that the signal arises from reciprocal connections across the dorsal tegmental nucleus (DTN) and lateral mammillary nuclei (LMN) (Taube, 2007; Cullen & Taube, 2017)(Fig. 1B). First, most attractor models contain both excitatory and inhibitory sets of connections, and previous anatomical studies have shown that there are excitatory projections from LMN → DTN and inhibitory projections from DTN → LMN (Groenewegen & van Dijk, 1984; Hayakawa & Zyo, 1990). These sets of connections make it an attractive candidate for a site where an attractor network might reside. Second, studies that interfere with the HD signal in brain areas upstream from the DTN/LMN have shown that cells in the ADN become bursty, although the burst activity is not tied to any particular direction. In contrast, interfering with brains areas either in LMN/DTN or in areas downstream from them do not lead to bursty behavior in ADN neurons or in neurons in other efferent brain areas (Valerio & Taube, 2016). Researchers have interpreted these findings to suggest that impairing brain areas afferent to the HD cell generation zone leave the attractor network intact, but unstable, causing the activity hill to drift unpredictably across different directions.

One key component of a HD attractor network are cells that are referred to as rotation cells. These cells are simultaneously sensitive to both HD and angular head velocity (AHV) and were first postulated by Skaggs et al. (1995; also see McNaughton et al., 1991) (Fig. 1A). Rotation cells are critical for shifting the activity hill to a new orientation during a head turn as they control which HD cells on the ring receive input to determine the new orientation. Currently, rotation cells have *not* been reported in rodents in areas where the HD signal is generated. While ‘pure’ AHV only cells have been identified in several brainstem areas afferent to the attractor network site, including the supragenual nucleus and nucleus prepositus, few, if any, display concurrent direction-specific firing (Graham et al., 2023). AHV cells alone, though, are not sufficient to shift the activity hill properly because they are connected to all the HD cells on the ring and would be unable to differentiate which HD cells need to receive the AHV information in order to accurately shift the activity hill to a new orientation. Downstream from LMN, HD cell firing in the ADN is mildly correlated with AHV (Taube, 1995; Taube & Muller, 1998); however, plots of firing rate vs. AHV for these cells showed little AHV tuning (Clark et al., 2024). Within the LMN – DTN network, ‘pure’ AHV cells (i.e., cells that are not conjunctive with HD) have been reported in both LMN and DTN, but for the most part did not display any directional selectivity (Stackman & Taube, 1998; Bassett et al., 2001; Sharp et al., 2001b). Further, similar to ADN HD cells, LMN HD cells were positively correlated with AHV, but AHV tuning curves were not constructed for these cells (Blair et al., 1998; Stackman & Taube, 1998). In the DTN, only a few HD cells have been reported, and except for one cell, they did not display any AHV sensitivity (Bassett & Taube, 2001; Sharp et al., 2001b).

In sum, both pure (classic) HD cells and pure AHV cells have been identified in a number of brain areas. While an attractor network remains the leading explanation for the generation of the HD signal in mammals, a key component of this model, the rotation cell, with strong tuning to both parameters (HD+AHV), has yet to be identified in mammals in areas where the HD signal is generated. Here we report the identification of such cells in both LMN and DTN. We show that there are two distinct types of HD cells in each of these brain areas – with one type displaying no AHV sensitivity while the second type exhibiting robust AHV tuning. We also demonstrate that a significant proportion of these conjunctive HD+AHV cells are also sensitive to the animal’s linear head velocity (speed).

## Results

A total of 74 HD cells were recorded from the LMN (Fig. 1C, top) while rats (n=27) foraged for small food pellets in a cylindrical enclosure (76 cm diameter) that were dispersed randomly around the apparatus. A single prominent visual cue attached to the inside wall of the cylinder served as the only intentional cue that the rats could use as a reference point. Recording sessions were 8-16 min in duration and were always conducted under lit conditions.

The HD cell population included a wide range of directionally tuned cells. Some cells displayed ‘classic’ tuning curves (e.g., Column 1: Figs. 2A, 3B,C, 4C, 5C), while others are best described as HD-modulated cells and contained broader directional firing ranges, higher background firing rates, and lower Rayleigh values (e.g., Column 1: Figs. 2C, 3E, 4B,D, 5A,D). In general, the LMN tuning curves, including those classified as classic, were not as sharp as those observed in the ADN (Taube, 1995), with broader directional firing ranges, with broader directional firing ranges. Nonetheless, all cells had significant Raleigh values (*p* < 0.01) and each cell passed a shuffle test (falling within the top 1^st^ percentile, see Methods). The inclusion of HD-modulated cells in the data set resulted in a lower mean Rayleigh value across this population, 0.516 ± 0.023 (range: 0.131 - 0.943, median: 0.512) compared to values from previous studies (ADN: 0.710 ± 0.018, median: 0.757, range: 0.330 - 0.954; PoS: 0.628 ± 0.020, median: 0.636, range: 0.305 - 0.973; Clark et al., 2024). Similarly, the mean directional firing range was larger than that reported in earlier studies, mean: 184.3 ± 5.5° (range: 70.9 – 291.1°, median: 190.6°). All recorded cells demonstrated stability throughout the recording session; the mean correlation between the first and second halves of a session were > 0.5 for 73 out 74 cells (mean: 0.872 ± 0.014, median: 0.915), with the remaining cell having a half session correlation of 0.399. As with previous studies, peak firing rates varied considerably among LMN HD cells, ranging from 4.9 to 239.3 spikes/s (mean: 56.6 ± 5.6 spikes/s; median: 40.6 spikes/s). Supplementary Table 1 summarizes HD cell properties among LMN cells classified as HD cells.

**Figure 2.**
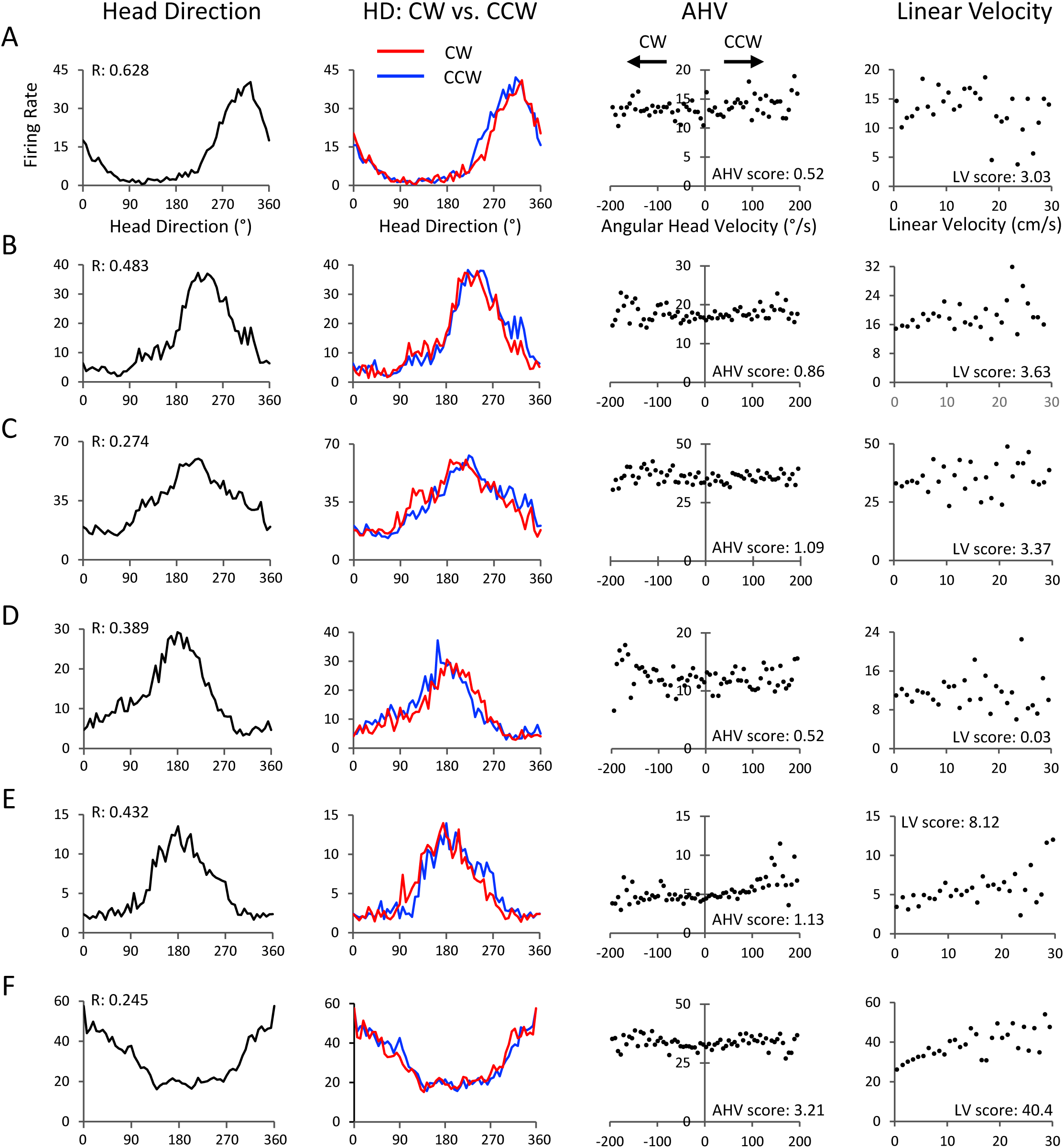
LMN AHV-independent cells. **A-F)** Six LMN HD cells are depicted that were not AHV sensitive. Each row is one cell. Starting at the left, the first column shows the HD x firing rate tuning curve. The second column depicts the HD tuning curve for the same cell but is divided into two functions based on whether the animal was turning CW (red) or CCW (blue). The third column shows the AHV tuning curve. The fourth column displays the linear velocity tuning curve. All axes are labelled as shown in the top row.

**Figure 3.**
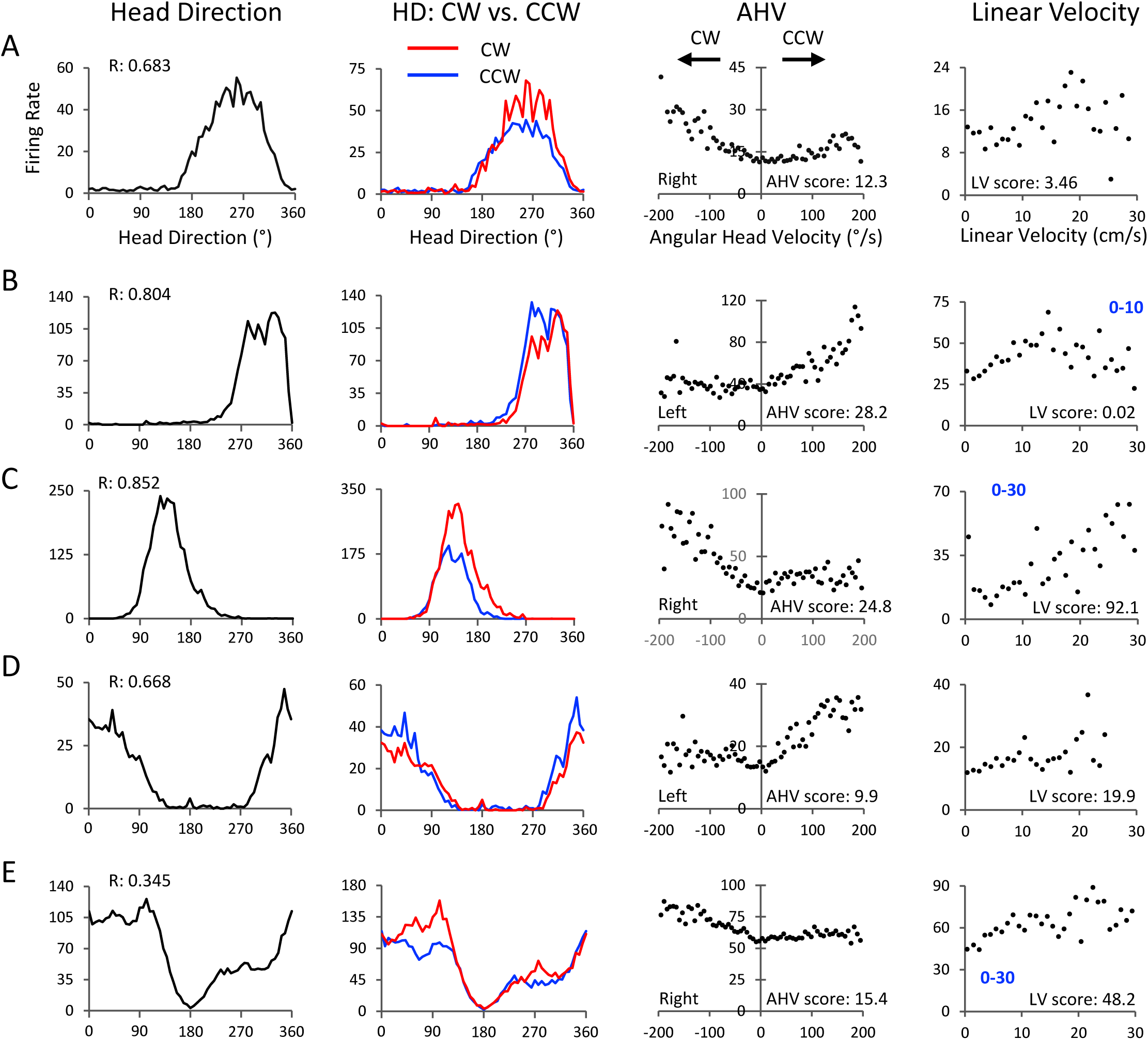
LMN Asymmetric-unresponsive HD+AHV cells. **A-E)** Five LMN HD cells are shown that were AHV sensitive and classified as Asymmetric-unresponsive cells. Each row is one cell. The cells in *A*, *C*, and *E* were recorded from the right hemisphere, while the cells in *B* and *D* were recorded from the left hemisphere. Note that the direction of AHV sensitivity (CW or CCW) is towards the hemispheric side where the electrode was implanted. Formatted columns are the same as Figure 2. All axes are labelled as shown in the top row.

**Figure 4.**
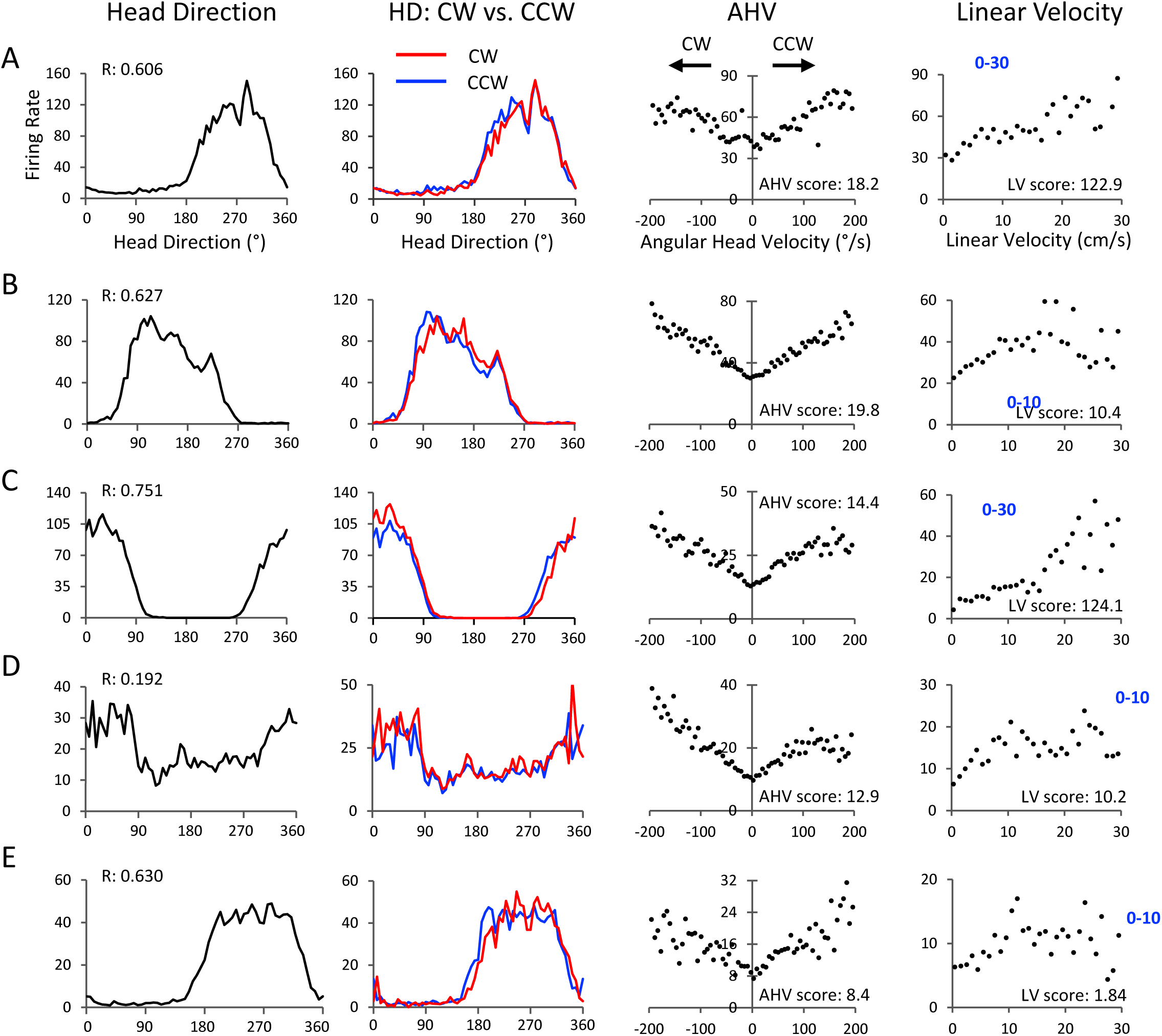
LMN Symmetric HD+AHV cells. **A-E)** Five LMN HD cells are shown that were AHV sensitive and classified as Symmetric cells. Each row is one cell. Formatted columns are the same as Figure 2. All axes are labelled as shown in the top row.

**Figure 5.**
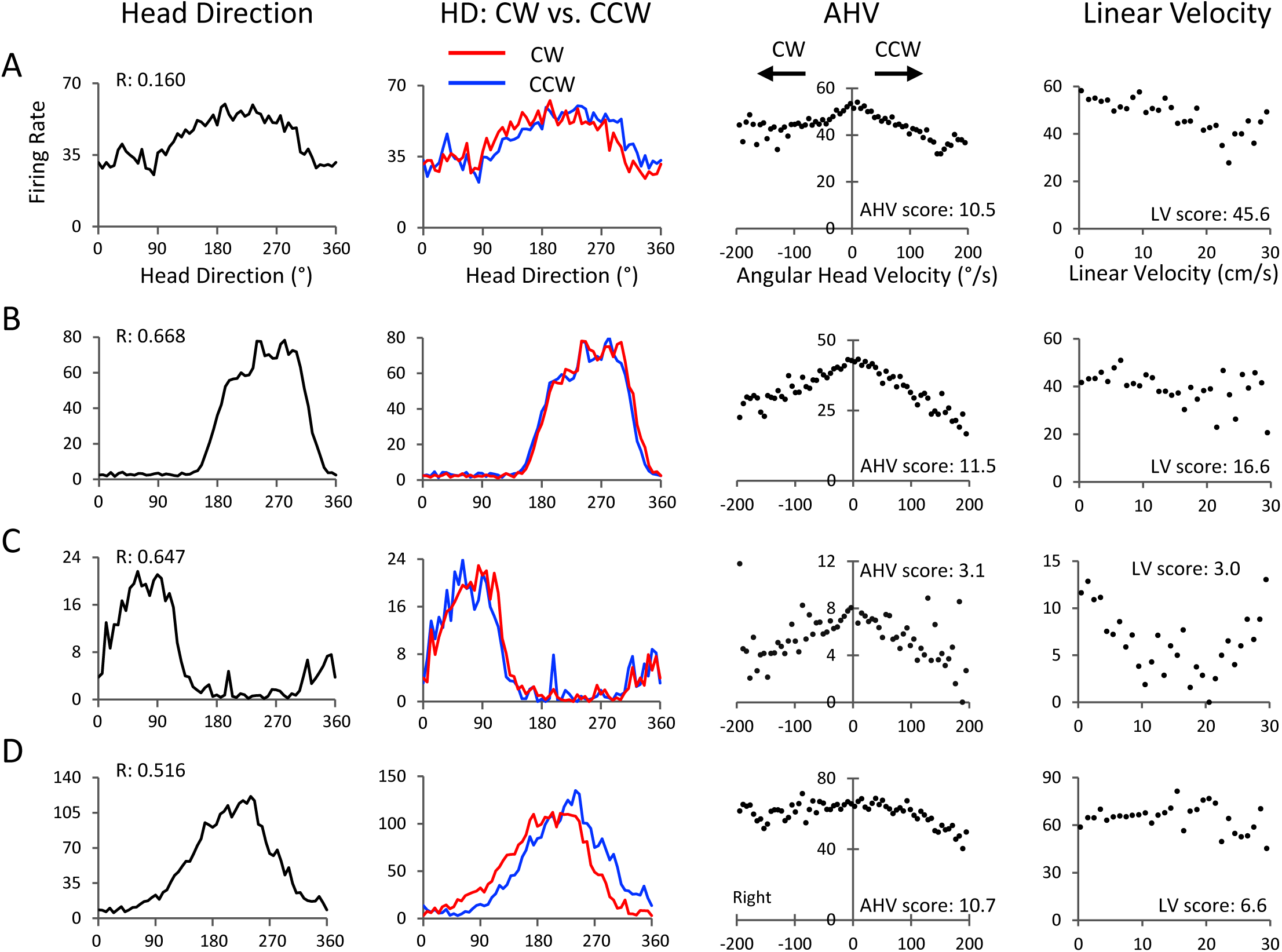
LMN Inverted HD+AHV cells. **A-D)** Four LMN HD cells are shown that were AHV sensitive and classified as Inverted cells. Each row is one cell. Formatted columns are the same as Figure 2. All axes are labelled as shown in the top row.

We next determined whether these HD cells were also sensitive to the animal’s AHV. 27 of the 74 LMN HD cells (36.5%) passed our AHV criteria test (see Methods) and were classified as AHV-dependent. The remaining 47 cells (63.5%) showed no significant AHV sensitivity in their tuning curves and were considered AHV-independent. Column 3 of Figures 2-5 show AHV tuning plots for 6 cells that were not tuned to AHV (Fig. 2) and for 14 cells that were classified as AHV dependent (Figs. 3-5). Across the AHV-dependent cells there was a wide range of AHV sensitivity from mild (e.g., Figs. 4A, 5C) to strong (e.g., Figs. 3C,E, 4B,D, 5B).

To measure the strength of AHV sensitivity we calculated an AHV score for each cell (see Methods). AHV-independent HD cells had lower AHV scores with 38 out of 47 cells having scores < 4.0. In general, AHV-independent HD cells that had higher AHV scores > 4.0 had poor looking AHV tuning curves and either did not pass the shuffle test or had a poor correlation between the first and second halves of the session and were therefore not classified as AHV-dependent (e.g., Fig. 2F). The mean AHV scores for AHV-dependent and AHV-independent HD cells were 12.48 ± 1.31 and 2.91 ± 0.51, respectively, and were significantly different from one another (F_(3,107)_ = 27.6, p < 0.00001; HSD = 9.57, *p* < 0.00001).

AHV tuning plots were also constructed for each cell using only samples that fell within the cell’s directional firing range (± 60° of the cell’s preferred firing direction). In general, the HD-independent and HD-dependent AHV plots were similar to one another (19 out of 27 cells), although the overall firing rates were higher for the HD-dependent plots because of the increased firing rate that occurred around the cell’s preferred firing direction compared to outside of it. Supplementary Figure 1 shows representative examples of the similarity between HD-independent and HD-dependent AHV tuning curves (Supplementary Fig. 1A,B) along with two examples when they were different (Supplementary Fig. 1C,D).

The AHV-dependent cells were further classified based on the shape of its AHV tuning curve. Previous studies on AHV cells have categorized them based on four subtypes (see Bassett & Taube, 2001; Graham et al., 2023):

1) Symmetric: cells that were AHV sensitive for both CW and CCW head turns.
2) Asymmetric: cells that were also sensitive for both CW and CCW head turns, but firing increased for one direction (e.g., CW) and decreased for the other direction (e.g., CCW).
3) Asymmetric-unresponsive: cells displayed increased firing for one turn direction, but were unresponsive in the opposite turn direction (flat tuning curve).
4) Inverted: cells that displayed decreasing firing rates for both CW and CCW head turns, with occasional cells showing a decreased firing rate in only one turn direction.

Using the HD-independent AHV plots from the 27 AHV-dependent cells, none of the cells were classified as purely asymmetric, 14 cells were classified as Asymmetric-unresponsive (Fig. 3), 8 cells were classified as Symmetric (Fig. 4), and 5 cells were classified as Inverted (Fig. 5) including one cell that was asymmetric in one direction and unresponsive in the opposite direction (Fig. 5D).

It is noteworthy that Asymmetric-unresponsive HD cells had unequal peak firing rate amplitudes in their CW vs. CCW HD tuning curves. This pattern is evident for all cells shown in column 2 of Figure 3 where the side that was more sensitive to AHV had a higher firing rate in the CW-CCW HD plot. In contrast, this observation was not evident for Symmetric or Inverted cells (compare the CW-CCW plots across Figs. 3-5, column 2).

We next examined whether the AHV sensitivity of Asymmetric-unresponsive cells was related to the hemisphere from which the cells were recorded. In the majority of cases (11 out of 15, 73.3%) the responsive portion of the AHV tuning curve corresponded to head turns toward the same side as the implanted electrode array (Fig. 3A-D). However, there were four cells (26.7%), all recorded from one rat, that showed the opposite response pattern (Supplementary Fig. 2). Indeed, all Asymmetric cells recorded from this rat were AHV sensitive for head turns opposite the implant side, differing from the response patterns observed in other rats (n=8). The reason for this difference is not clear.

We next analyzed whether the conjunctive HD-AHV cells correlated with any particular HD cell properties compared to HD cells that were not AHV sensitive. We found that conjunctive HD+AHV cells were more likely to have higher peak firing rates compared to AHV-independent HD cells (Fig. 6A left; F_(3,107)_ = 12.41, p < 0.00001; HSD= 43.72, *p* < 0.001). Furthermore, within the AHV-dependent HD cell population, a strong correlation was observed between a cell’s peak firing rate and its AHV score (*r*_(25)_ = 0.794, *p* < 0.01) (Fig. 6B). However, there were no significant differences between AHV-dependent and AHV-independent HD cells in measures of Rayleigh values, directional firing range, directional information content, or half-session correlation (a measure of spatial stability across the session); all *p’s* n.s. Table 1 summarizes these comparisons for LMN cells. Further, there was no propensity for higher AHV scores to be associated with a particular AHV cell type (i.e., symmetric vs. asymmetric-unresponsive vs. inverted).

**Figure 6.**
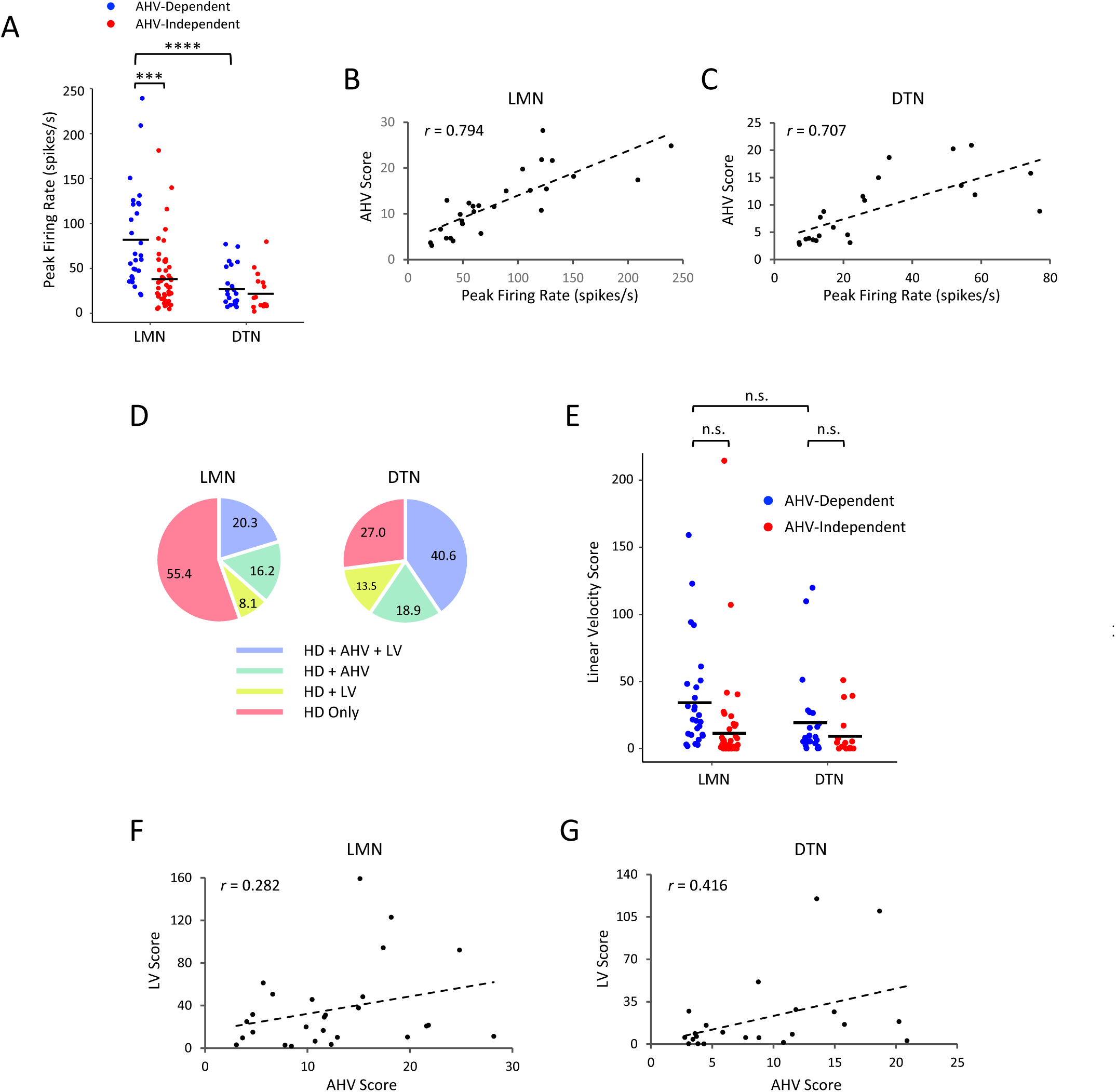
LMN and DTN properties. **A**) Peak firing rate for all AHV-dependent and AHV-independent HD cells in LMN (left) and DTN (right). Overall, HD cells that were AHV-dependent had significantly higher peak firing rates than AHV-independent cells for LMN, but not DTN. *** *p* < 0.001, \*\*\*\**p* < 0.0001. **B**) Relationship between a cell’s peak firing rate and its AHV score for all LMN AHV-dependent HD cells. Cells with higher peak firing rates generally had larger AHV scores. Dashed line shows the best-fit line for the data. **C**) Same as *B* except for DTN AHV-dependent HD cells. **D**) Pie charts showing the proportion of cell types in LMN (left) and DTN (right). Note, only HD cells are shown. The proportion of AHV-dependent cells and those with LV sensitivity were higher in DTN than in LMN. **E**) LV scores for the AHV-dependent and AHV-independent populations for LMN (left) and DTN (right). There was no significant difference between LV scores for AHV-independent and AHV-dependent cells for both LMN and DTN, as well as between the two brain areas. **F,G**) Plots displaying the relationship between AHV vs. LV scores across all AHV dependent HD cells (HD+AHV) for LMN (*F*) and DTN (*G*). Dashed lines indicate the best-fit line for the data.

**Table 1.**
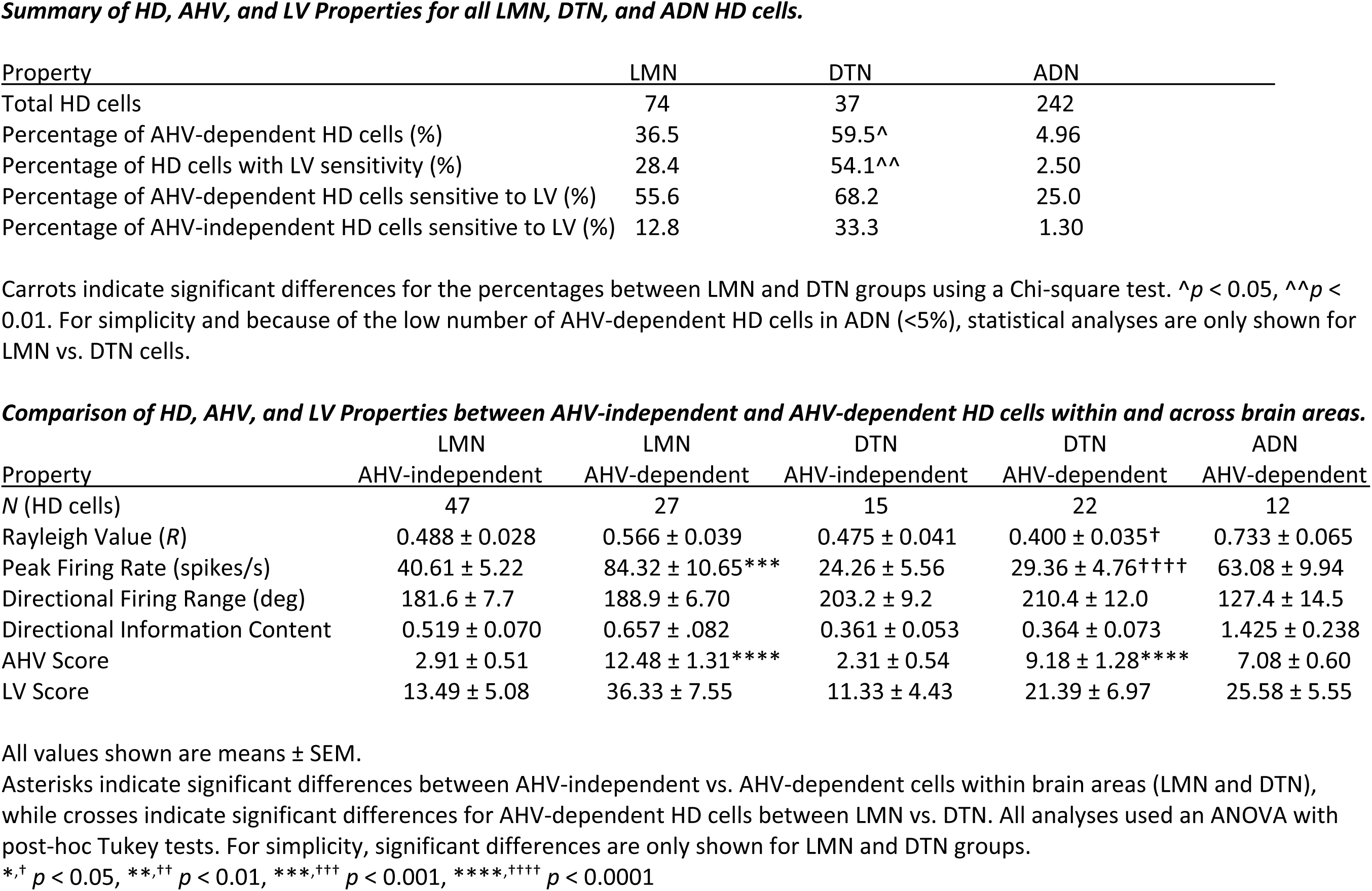

In summary, two distinct types of HD cells were identified in the LMN, with one type sensitive to the animal’s AHV, while the second type was not sensitive to the animal’s AHV.

### Sensitivity to linear velocity (speed)

We next assessed whether the HD cells were sensitive to linear head velocity (LV) by analyzing their firing rate vs. LV tuning plots (see Methods). These plots were constructed using data that excluded samples when the rat was turning its head at speeds > 10 °/sec as this would confound the analysis for LV. All HD cells depicted in Figures 2-5 show their LV tuning curve in column 4. While some cells display LV sensitivity throughout their tuning curve (i.e., 0-30 cm/s)(Figs. 3C, 4A,C, 5A), some cells only contained LV sensitivity in the range of 0-10 cm/s (Figs. 3B,E, 4B,D). For purposes of describing how many cells contained LV sensitivity at the population level, we grouped both the 0-30 cm/s and the 0-10 cm/s groups together. Of the 74 HD cells, 21 cells (28.4%) were classified as sensitive to LV, including 6 cells that were only LV sensitive in the 0-10 cm/sec range. Of these 21 LV sensitive HD cells, 15 of them (71.4%) were also sensitive to AHV. Further, of the 27 AHV-dependent HD cells discussed above, 15 of them (55.6%) were sensitive to LV. Figure 6D left is a pie chart summarizing the percentage of HD cells in each category. Figure 6E left plots all HD cells based on their LV score and whether they were classified as AHV-dependent or AHV-independent. Overall, AHV-dependent cells had higher LV scores than AHV-independent cells (means: 36.33 ± 7.55 vs. 13.49 ± 5.08), but the difference was not significant (F_(3,107)_ = 3.01, *p* < 0.05; HSD = 22.84, *n.s.*).

Cells that were sensitive to LV could either have positive slopes (increased firing with increasing LV), n = 19 (Figs. 3B,C,E, 4A-D), or negative slopes (deceased firing with increasing LV), n=2 (Fig. 5A). LV sensitive cells were found across all three AHV-dependent cell types: Symmetric (n=7), Asymmetric-unresponsive (n=7), and Inverted (n=1). Thus, there was no apparent trend linking specific AHV cell subtypes with LV sensitivity among cells that were tuned to both AHV and LV. Next, we examined the AHV-independent population of HD cells (n=47). Among this group, only 6 cells (12.8%) were sensitive to LV (e.g., Fig. 2F). Finally, we examined the relationship between AHV and LV scores across all HD cells. Figure 6F left plots a cell’s AHV score versus its LV score for HD AHV-dependent cells; there was only a weak correlation between these two parameters (*r*_(25)_ = 0.282, *n.s.*).

### Dorsal Tegmental Nuclei

In a different set of animals (n=10), we recorded 37 HD cells from the DTN (Fig. 1C, bottom). The lower number of HD cells recorded in DTN compared to LMN reflects the much smaller percentage of HD cells present in the DTN compared to pure AHV cells (Bassett & Taube, 2001; Sharp et al., 2001). Many of the HD cells recorded in the DTN were better described as HD-modulated cells, as the cells had broad directional firing ranges and lacked sharp tuning curves (see Fig. 7A,D,F,H, column 1). Supplementary Table 1 summarizes the mean values for different HD cell properties. Compared to LMN HD cells, DTN cells had significantly lower mean peak firing rates, Rayleigh values, and information content scores, while their directional firing ranges were significantly broader.

**Figure 7.**
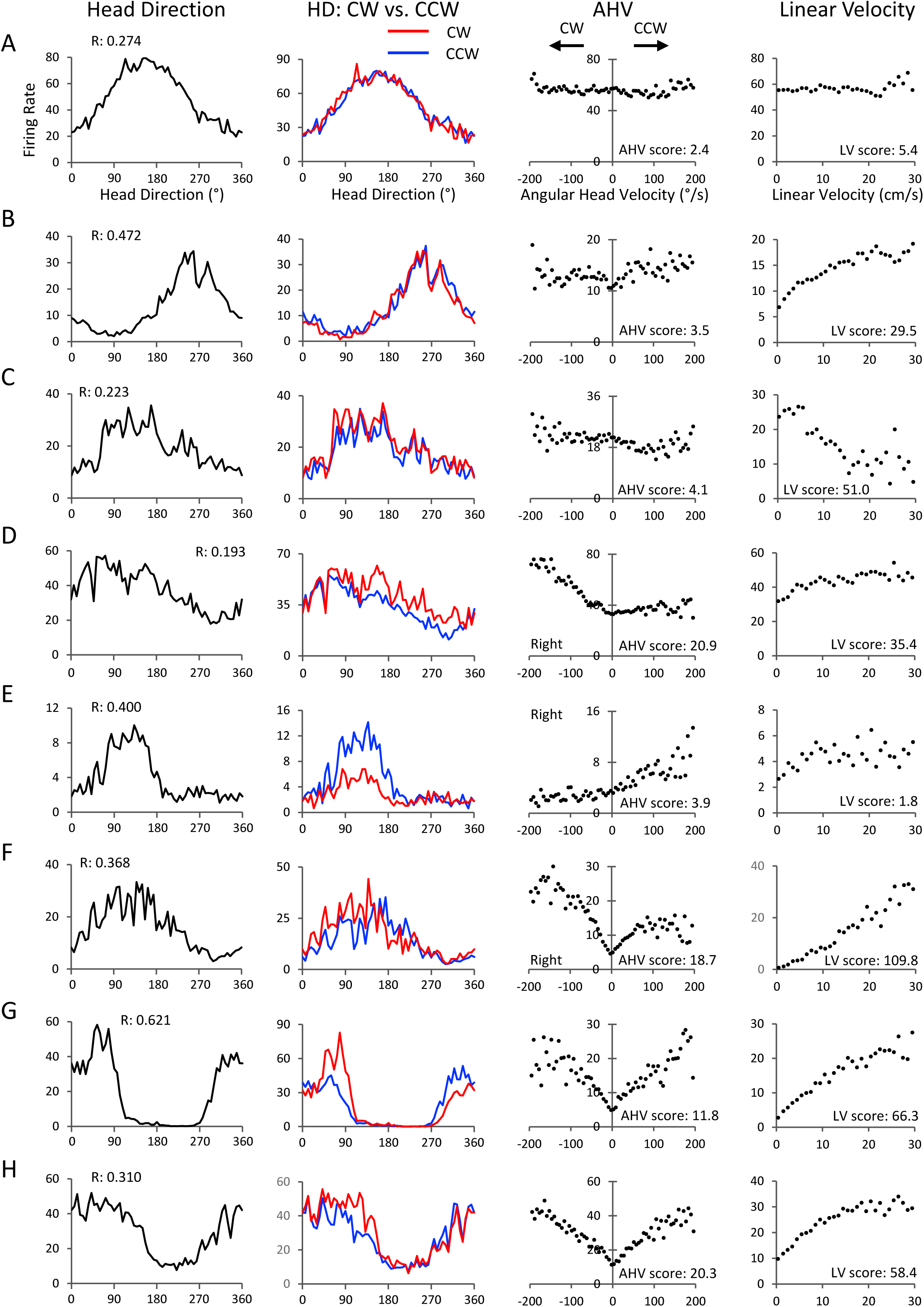
DTN HD cells. Eight DTN HD cells are shown with each row representing one cell. **A-C)** Three HD cells that were not sensitive to AHV. The cell in *A* was not sensitive to LV, while the cell in *B* was sensitive to LV. The non-AHV HD cell in *C* is notable because it has a negative sloped tuning curve for linear velocity. **D-F)** Three HD+AHV cells that were sensitive to AHV and classified as Asymmetric. The cells in *D* and *E* were recorded from the same animal. Note that the cell in *D* is sensitive for CW directions, while the cell in *E* is sensitive for CCW directions. Thus, cells sensitive to different turn directions can be found within the same DTN nucleus. While the cell in *D* appears mildly sensitive to LV (LV score: 35.4), it was not classified as such because the half session analysis indicated the LV sensitivity was not present in the second half of the session. The cell in *E* was also not classified as sensitive to LV (LV score: 1.8). The cell in *F* was symmetric at low AHVs (0-70°/s), but asymmetric at high AHVs (>70°/s). **G, H)** Two HD+AHV cells that were sensitive to AHV and classified as Symmetric. The cells in *G* and *H* were both sensitive to LV. Formatted columns are the same as Figure 2. All axes are labelled as shown in the top row.

Using the same criteria to classify cells as AHV sensitive, 22 of the 37 DTN HD cells (59.5%) were classified as sensitive to AHV, with 9 cells classified as asymmetric-unresponsive (Fig. 7D-F, third column), 13 cells classified as symmetric (Fig. 7G-H, third column), and no cells classified as inverted. This proportion of AHV-dependent cells was higher and significantly different from the percentage in LMN (chi-square = 5.28, *p* < 0.05.). The remaining 15 cells (40.5%) showed no AHV sensitivity in their tuning curves and were classified as AHV-independent (Fig. 7A-C, third column). In contrast to LMN, there was no difference in the peak firing rates between AHV-dependent and AHV-independent cells in DTN (Fig. 6A, right). However, as with LMN, AHV scores for DTN AHV-dependent HD cells correlated strongly with peak firing rate (*r*_(20)_ = 0.707, *p* < 0.01; Fig. 6C). The mean AHV score for AHV-dependent DTN cells was 9.18 ± 1.28 and was significantly different from AHV-independent DTN cells (F_(3,107)_ = 27.57, p < 0.00001; HSD = 6.87, *p* < 0.0001), but not significantly different from AHV scores from LMN AHV-dependent cells (12.48) (F_(3,107)_ = 27.57, *p* < 0.00001; HSD = 3.30, *n.s.*). When examining the HD+AHV asymmetric population in DTN, we found that cells sensitive to different turn directions (CW vs. CCW) could be found within the same DTN nucleus. For example, the two cells depicted in Figure 7D,E were recorded from the same animal in the same hemisphere. The cell in D was sensitive for CW directions, while the cell in E was sensitive for CCW directions.

In terms of sensitivity to LV, 20 of 37 DTN HD cells (54.1%) were also sensitive to LV. Among the 22 AHV-dependent cells, 15 of them (68.2%) were also sensitive to LV. Thus, the percentage of LV sensitive cells in DTN was greater than in LMN (54.1 vs. 28.4%, chi-square = 6.98, *p* < 0.01), but the percentage of LV sensitivity amongst AHV-dependent cells was similar (68.2 vs. 55.6%, chi-square = 0.815, *n.s.*) between the two areas. Similar to LMN cells, for some DTN cells (n=6) the LV sensitivity was only present in the 0-10 cm/sec range (Fig. 7D,E). All DTN sensitive LV cells had positive correlations between LV and firing rate except for one cell that had a negative correlation (Fig. 7C); this cell was not sensitive to AHV. For DTN HD cells, LV scores were not significantly different between AHV-dependent and AHV-independent groups (F_(3,107)_ = 3.01, p < 0.05; HSD = 10.06, *n.s.*; Fig. 6E, right). We then examined the AHV-independent population of HD cells (n=15). Within this group, only 5 cells (33.3%) were sensitive to LV (Fig. 7B,C). Figure 6D right shows the proportion of different cell types found for DTN HD cells.

Finally, we investigated the relationship between AHV and LV scores across AHV-dependent cells (n=22; Fig. 6G right). We found a weak positive correlation between these two parameters, but it was not significant (*r*_(20)_ = 0.416, *n.s.*). Table 1 summarizes the differences between AHV-independent and AHV-dependent HD cells for DTN, as well as comparing them to LMN HD cells.

### Comparison to HD cells in the ADN Thalamus

Previous studies have shown that the ADN thalamus contains a substantial population of HD cells, with a high percentage of them classified as classical ones. Further, the ADN thalamus is thought to function as a key node in broadcasting the HD signal to several cortical structures. To assess the extent of conjunctive tuning between direction-specific firing, AHV, and LV in the ADN, we analyzed a population of ADN HD cells recorded in previous studies (n=242). Using the same AHV criteria as described above for LMN HD cells, we found that only a small number of ADN HD cells, n=12 (4.96%), were conjunctively tuned to AHV. Among this group, 9 cells were classified as symmetric, 1 cell was classified as asymmetric-unresponsive, and 2 cells were classified as inverted (examples of a Symmetric cell and an Inverted cell are depicted in Fig. 7 of Clark et al., 2024). This proportion of AHV-dependent HD cells in the ADN is much less than in the LMN and DTN. Further, even among the cells that were classified as AHV-dependent in ADN, the sensitivity to AHV in this population was significantly weaker compared to LMN (but not DTN), as indicated by the lower mean AHV score for this group (7.08 ± 0.60) compared to 12.48 ± 1.31 for LMN AHV-dependent cells (F_(2,58)_ = 4.06, p < 0.05; HSD = 5.40, *p* < 0.05).

In terms of LV sensitivity, we found that only 6 out of 242 cells (2.5%) were classified as sensitive to LV, which is lower than the percentage that was sensitive to AHV (4.9%), and much lower than the proportions for LMN (44.6%) and DTN (48.6%). Of the 12 AHV-dependent ADN HD cells, 3 of them (25.0%) were also sensitive to LV.

## Discussion

Our results demonstrate that there are two distinct types of HD cells in the LMN – one that is conjunctively dependent with AHV and one that is AHV-independent. About one third of the HD cells in LMN were sensitive to AHV and exhibited one of three different types of AHV tuning - Symmetric, Asymmetric-unresponsive, and Inverted. These AHV-dependent HD cells form a separate population of cells from pure AHV cells, which are not sensitive to HD, and have also been reported in the LMN (Stackman & Taube, 1998). A similar pattern was observed in the DTN, where an even larger proportion of HD cells were AHV dependent (∼60%). As with the LMN, this HD AHV-dependent population is different from the pure AHV, non-HD population of cells previously reported in DTN (Bassett & Taube, 2001; Sharp et al., 2001b). In addition, the same three types of AHV-dependent cells were observed in DTN with comparable proportions as in LMN. Further, we did not find differences in HD cell properties between AHV-dependent and AHV-independent HD cells, although LMN HD cells that were AHV-dependent were more likely to contain higher peak firing rates.

In contrast, the ADN thalamus, an area that contains a high percentage of HD cells (Taube, 1995), contained very few AHV-dependent HD cells (< 5%), and even the ones that were identified as conjunctive, their AHV sensitivity was notably weaker than in LMN (Table 1). Earlier studies on ADN HD cells found a significant correlation with between cell firing and AHV (Taube, 1995; Taube & Muller, 1998). However, those analyses were based on correlating the spike time series with the AHV time series for all samples rather than AHV tuning curves. As reported here, when such tuning curves were constructed, very few ADN HD cells displayed AHV sensitivity that met our criteria.

The ADN receives a major input from LMN and is critical for transmitting the HD cell signal to various cortical areas including the postsubiculum, medial entorhinal, postrhinal, and medial precentral cortices (Goodridge & Taube, 1998; Winter et al., 2015; Mehlman et al., 2019; LaChance & Taube, 2024). Interestingly, HD cell recordings from most of these areas displayed little AHV sensitivity based on the construction of AHV tuning curves (Clark et al., 2024). In contrast, a recent study in mouse retrosplenial cortex identified both pure HD and conjunctive HD+AHV cells, with 71% being of the conjunctive type (Keshavarzi et al., 2022). They also identified pure AHV cells that were HD independent and included all three types of AHV tuning (symmetric, asymmetric, and inverted). Although yet to be demonstrated experimentally, it is likely that the RSC receives its HD information from the ADN directly, or indirectly from ADN → postsubiculum → RSC. In contrast, the AHV information encoded in RSC cells is unlikely to be derived from ADN since so few ADN HD cells (<5%) encode a significant amount of AHV information. Nonetheless, we cannot exclude the possibility that the small number of ADN HD+AHV cells that are present are sufficient to provide AHV information to RSC. Although the RSC AHV information could be self-generated, it is more likely that the RSC receives AHV information from other cortical (or subcortical) areas as Hennestad et al. (2021) found that a vestibular-dependent AHV signal is widespread across many cortical areas. Consistent with this view, when lesions are made to ADN, HD cells in the striatum and medial precentral cortex lose their direction-specific firing, but AHV cells in the same locale are unaffected and maintain their AHV sensitivity (Mehlman et al., 2019) - demonstrating that the ADN is not the primary source of AHV information to these areas. Taken together, our findings highlight the distinctions between HD signaling in subcortical regions (LMN and DTN) and the ADN, emphasizing the importance of the LMN-DTN network in generating the conjunctive HD+AHV signals and the more specialized role of ADN in conveying HD information to cortical structures.

### Sensitivity to linear velocity (LV)

Our results also showed that a large number of HD cells, in both LMN and DTN, were also sensitive to LV – especially among those cells that were also sensitive to AHV. This finding was somewhat unexpected, but it does not appear to be an artifact of our methodology in calculating LV, since applying an alternative method for calculating LV that used longer sampling intervals (250 msec episodes rather than 16.6 msec intervals) returned similar results (data not shown). Given that angular and linear velocity encode different parameters of movement, the functional significance of these conjunctive HD+AHV+LV cells is not clear, although it is possible that these cells signal the extent of movement in a particular HD. Because our animals were moving in an open-field environment, we were limited in the number of samples collected when the rat was moving forward linearly, even when we restricted samples to data points where the AHV component was < 10°/sec. Recording these cells when an animal is performing a task on a linear track, which would limit the extent of head turns, would aid in addressing the extent of sensitivity to LV.

### Attractor networks and rotation cells

A major goal of this study was to determine whether ‘Rotation cells’ exist within the brain areas believed to generate the HD signal. McNaughton and colleagues (McNaughton et al., 1991; Skaggs et al., 1995) originally proposed that cells sensitive to both HD and AHV were necessary for an attractor network to effectively update the HD signal following a head turn. Pure AHV cells, which arise from the vestibular nuclei and convey information about AHV to the HD network (Graham et al., 2023), are not sufficient by themselves to update the HD signal because if they were connected directly to all HD cells, they would be unable to differentiate which HD cells need to be activated in order to shift the activity hill correctly. Thus, there must be a mechanism whereby only the cells currently participating in the activity hill are to be excited by AHV in the appropriate direction (CW or CCW). For this event to occur, Skaggs and McNaughton postulated the existence of two additional rings of so-called rotation cells – one ring responsive to CW turns and the other ring responsive to CCW turns. The rotation cells in each ring received inputs from both pure HD cells in the HD ring and from pure AHV cells tuned to either CW or CCW directions (Fig. 1A). The rotation cells, in turn, were selectively activate the HD ring, to move the activity hill in the appropriate CW or CCW direction. This mechanism ensures accurate rotation of the HD activity hill during head turns (see Fig. 1). Thus, rotation cells are critical components of a properly functioning HD attractor network.

Previous studies have reported rotation-like cells, but only in cortical areas, and not in regions where the HD signal is believed to be generated (e.g., Wilber et al., 2014; Keshavarzi et al., 2022). In LMN and DTN, almost all identified cells have either been pure HD or pure AHV cells and only a single conjunctive HD+AHV cell has been reported (e.g., see Fig. 3 of Sharp et al., 2001). Here, we show that significant proportions of rotation cells are present in both LMN and DTN. Clearly, not all HD cells are conjunctively sensitive to AHV, as both LMN and DTN also contain significant proportions of pure HD cells that are not AHV sensitive. Thus, the HD+AHV cells represent a distinct subset. Importantly, in the attractor network(s) proposed by Skaggs and McNaughton, there was only one type of rotation cell – one that was tuned across a wide range for both CW and CCW turns. These rotation cells were hypothesized to receive inputs from vestibular-sensitive AHV cells, which (in monkeys) were known to fire in an asymmetric manner throughout the CW-CCW range (Fuchs & Kim, 1975; Gdowski & McCrea, 1999). We found, however, that there was an absence of this asymmetric cell type, and instead, we identified three other distinct types of HD+AHV cells: symmetric, asymmetric-unresponsive, and inverted. Although similar to a ‘pure’ asymmetric cell type, the asymmetric-unresponsive cell was only AHV sensitive in one turn direction - usually ipsilateral to the recording site (i.e., CW AHV sensitivity corresponded to right hemisphere recordings and vice versa). This pattern, however, was reversed in one rat, indicating potential variability across individuals.

While both brain areas contained the same types of conjunctive HD+AHV cells, the proportions of different cell types differed slightly between LMN and DTN. For instance, DTN had a slightly higher proportion of symmetric cells than LMN, while LMN had more asymmetric-unresponsive cells. Notably, inverted HD+AHV cells were absent from the DTN. Nonetheless, the sizeable proportion of symmetric type HD+AHV cells highlights the importance of this cell type and raises the issue of what function they might serve. While the function of the asymmetric type presumably serves to push the activity hill of the attractor network in the direction of the head turn, the function(s) of both the symmetric and inverted types are less clear. One possibility is that the asymmetric type cell is not sufficient to overcome the stability of the network to move the activity hill to a new state in the face of a head turn and that a further boost of excitation is required. This boost could be accomplished by the symmetric type cells, which would serve to aid in moving the activity hill in either direction away from a stable state, while the asymmetric cells would serve in controlling the direction the activity hill needs to move. Alternatively, the symmetric AHV cells may be used to calibrate or fine-tune the HD system for preventing drift of the HD activity hill, as the initial connections following development are unlikely to be balanced perfectly (Stratton et al., 2010). In terms of the inverted conjunctive cells, they may serve to suppress any possible excitation in the network for HDs that are distant to the current HD or they could provide a brake on the system to keep the activity hill from moving continually.

Our findings support the hypothesis that a set of attractor rings, originally proposed by Skaggs et al. (1995), form the basis for the generation of the HD signal. However, their model (Fig. 1A) only contained one type of Rotation cell – one that would be classified as asymmetric throughout head turns in both directions. The only AHV cell type we observed similar to this firing pattern was the asymmetric-unresponsive one where AHV sensitivity was only present for one turn direction and not sensitive to AHV in the opposite direction. Further, as discussed above, the attractor model will also require the addition of other HD+AHV cell types that include both symmetric and inverted firing patterns.

Recent work in the *Drosophila* central complex have identified circuits involved in encoding directional heading (Seelig et al., 2015; Hulse and Jayaraman, 2020) and some of these studies have found HD+AHV conjunctive type cells – similar to the rotation cells postulated by Skaggs and McNaughton. Such cells, referred to as P-EN cells have been identified in the ellipsoid body of *Drosophila* (Green et al., 2017; Turner-Evans, 2017). Most of the data reported have focused on asymmetric type cells, and it is not clear if there is a substantial proportion of symmetric type cells similar to the ones reported here in rodents. Moreover, to our knowledge, no inverted type rotation cells have been identified. Thus, while parallels between HD systems in insects and mammals are emerging, it remains to be determined whether the underlying neural mechanisms contributing to the HD signal in *Drosophila* are ultimately going to be similar to those in a mammalian system.

### Conclusions

We have demonstrated that there are two fundamentally distinct types of HD cells in LMN and DTN – brainstem areas where the HD signal is believed to be generated. One type is AHV-dependent (referred to as rotation cells), while the second type is *not* sensitive to AHV. In addition, both cell types were frequently found to be sensitive to the animal’s linear head velocity – particularly in the DTN. In addition, both cell types were frequently found to be sensitive to the animal’s linear velocity, further expanding their functional profile. Ring attractor networks, hypothesized to generate and update the HD signal, require AHV-dependent HD cells to properly shift the activity hill in response to head turns. Our identification of this requisite cell type in LMN and DTN, along with the presence of pure HD and pure AHV cells in these same brain areas, strongly supports the existence of an attractor-style network in these brainstem regions. Collectively, our results provide crucial evidence for the neural mechanisms underlying the generation and dynamic updating of the HD signal within the LMN and DTN circuit.

## Methods

### Subjects and Apparatus

Experiments were conducted using three separate groups of Long-Evans rats. The first group had electrodes implanted targeting the LMN, the second group had electrodes implanted to target the DTN, and the third group consisted of previously recorded rats with electrodes implanted into the ADN thalamus. Animals from all three groups were recorded in the same shaped apparatus – a gray cylinder that contained a single prominent landmark cue and was located in a 10’ x 10’ room. All procedures were conducted in accordance with an institutionally approved animal care protocol and in compliance with guidelines described in the *National Institutes of Health Guide for the Care and Use of Laboratory Animals*.

The LMN group included 27 female rats, weighing 250–300 g at the start of the experiment, 13 of which came from previous studies (encompassing 37 out of the 74 HD cells analyzed here; Stackman & Taube, 1998; Yoder et al., 2015). The DTN group consisted of 10 female rats of similar weight. All rats were maintained on a food-restricted diet of ∼15 g/day, and water was available ad libitum. The rats were housed individually in transparent plastic cages and kept on a 12:12-h light-dark cycle. Training and HD cell screening occurred during sessions in which the rats foraged for food pellets in a gray-colored cylinder (71 cm high, 76-cm diameter). A white sheet of cardboard attached to the inside wall of the cylinder covering ∼100° of arc served as the only intentional visual cue that could be used for orientation. The cylinder was placed on a sheet of gray photographic backdrop paper, which was replaced after each recording session to remove olfactory cues from the previous session. A black curtain (2.5 m in diameter) surrounded the cylinder and hung from the ceiling to the floor. Eight overhead DC lamps spaced uniformly in a circle provided room illumination. A color video camera (Sony XC-711, Tokyo, Japan) was attached to the ceiling directly above the cylinder. A sound machine was attached to the ceiling adjacent to the video camera and played white noise to help mask any external auditory cues. A ceiling-mounted food dispenser that was also positioned adjacent to the video camera automatically ejected sucrose pellets (20 mg; Dustless Precision Pellets, Bio-Serv, Flemington, NJ, USA) on a 30 sec random interval schedule.

### Training Procedures

Rats were first trained to forage for food pellets in the cylinder. They were removed from their cages and placed in the cylinder for 10 min. During training the rats were encouraged to forage for small food pellets dropped randomly into the cylinder from the automatic food pellet dispenser. The procedure was repeated daily for 5–7 days. Upon completion of training, rats engaged in nearly continuous pellet search behavior.

### Electrodes

Electrodes consisted of either a bundle of 10 single wires (n=16 for LMN; n=5 for DTN), an array of 8 stereotrodes (n=11 for LMN; n=3 for DTN), or an array of 4 tetrodes (n=2 for DTN). The single-wire electrodes were built as previously described (Kubie, 1984) and consisted of a bundle of ten 25 µm nichrome wires (California Fine Wire, Grover City, CA, USA) insulated except at the tips. The wires were threaded through a 26 gauge stainless steel cannula and connected to a modified Augat plug. Stereotrodes and tetrodes were made by spinning 17 µm nichrome wires into a compact bundle composed of either two (stereotrodes) or four wires (tetrodes; Winter et al., 2015). These wires were threaded through a smaller 32 gauge cannula that was connected to a modified Mill-Max connector. Before surgical implantation, the electrode bundle was sterilized and coated (except for the tips) with polyethylene glycol (Carbo wax) to provide stability to the electrode wires as they were positioned in the brain.

### Surgical Procedures

After training, each rat was anesthetized with either Nembutal (1.0 ml/kg, i.p.), a ketamine-xylazine mixture (2 ml/kg, i.m.), or isoflurane gas and placed into a stereotaxic frame (David Kopf Instruments, Tujunga, CA, USA). An electrode array was then implanted just above the targeted site. Electrode coordinates varied slightly between different animals but were within ± 0.1 mm of the following parameters with respect to bregma: LMN: anterior/posterior - 4.6 mm, medial/lateral ± 1.0 mm right or left, and ventral 8.1 mm from the cortical surface (Paxinos and Watson 1998); DTN: anterior/posterior -9.1 mm, medial/lateral ± 0.3 mm right or left, and ventral 5.4 mm from the cortical surface (Paxinos and Watson 1998). Four to eight stainless steel screws were placed in the skull plates, and dental cement securely anchored the electrode assembly in place. This assembly could be manipulated in the dorsal/ventral plane by turning three drives screws that were part of the electrode assembly. Rats were given the analgesic ketoprofen (3-5 mg/kg, i.m.) after the operation for two consecutive days and were given 1 week of recovery before commencing recording for HD cells.

### Recording & Video Tracking

One of two different recording systems was used for the experiments. For cells recorded with single wires, we used an analogue based system using window discriminators to isolate cell waveforms (LMN: 16 rats; DTN: 5 rats; see Taube, 1995 for a description of data acquisition). For cells recorded with stereotrodes or tetrodes, we used a NeuraLynx Digital system (Bozeman, MT, USA) (LMN: 11 rats; DTN: 5 rats; see Graham et al., 2023 for a description of data acquisition).

For both methods of recording, each rat’s microelectrodes were monitored for cellular activity while the rat foraged for food pellets in the cylinder. The rats were transported from the animal colony room into the screening room, and no attempt was made to disorient the rats before each session. For cells recorded from single wires, each of the 10 microelectrodes was monitored on an oscilloscope, and the rat’s directional heading was observed on a video screen via the overhead video camera. The electrode wires were advanced gradually over several weeks while screening for HD cells, in which their electrical waveform was of sufficient amplitude for isolation from background noise. For recordings from single wires, we monitored neural activity by passing the signals through a field-effect transistor (FET) that was in a source-follower configuration and then through an overhead commutator (Biela Idea Development, Anaheim, CA, USA) to an amplifier (Grass Instruments P5 Series, West Warwick, RI, USA). Signals were referenced to electrical activity recorded on a nearby wire and then differentially amplified (20,000), band-pass filtered (300 –10,000 Hz, 3 dB/octave; Peavey Electronics PME8, Meridian, MS, USA), and sent through a series of window discriminators (BAK Electronics model DDIS-1, Germantown, MD, USA). The final signal was then displayed on an oscilloscope (Tektronix model 2214, Beaverton, OR, USA). When the waveform of a HD cell could be isolated successfully, two light-emitting diodes (LEDs) were activated via the recording cable to monitor the animal’s HD via the overhead video camera. The headstage included a red LED positioned ∼1 cm posterior to the headstage and a green LED positioned ∼6-10 cm further away over the rat’s back. For single wire recordings, the x,y-coordinates of the LEDs were monitored at 60 Hz by an automated video-tracking system (Ebtronics, Brooklyn, NY, USA). During recording sessions the LED coordinates and the neural activity were acquired by a data acquisition interface board (National Instruments DIO-96, Austin, TX, USA) in a Macintosh computer.

Signals originating from the stereotrodes or tetrodes were preamplified by unity-gain operational amplifiers on an HS-27 headstage, then further amplified (NeuraLynx Digital Lynx SX) and then bandpass filtered (600–6000Hz) using an ERP-27 system (NeuraLynx). When the neural signals crossed a preset amplitude threshold (30–60 mV), they were time stamped and digitized at 32 kHz for 1 msec. The waveform characteristics were then analyzed off-line using SpikeSort 3D (NeuraLynx). The location and directional heading of the rat were also tracked in a similar manner as above using red and green LEDs.

For both recording systems, the cell’s firing rate relative to the animal’s HD was computed off-line with programs written with LabVIEW software (National Instruments, Austin, TX, USA). A cell’s activity was usually recorded for 16 min (some sessions were only 8 min long) while the rat foraged for food pellets inside the cylinder. If no HD cell activity was detected, the entire electrode array was advanced 25–50 µm ventrally and electrical activity was screened again 24 h later. Screening for HD cells generally occurred over the course of 2–3 months.

### Data Analysis

For each recording session, a firing rate vs. HD tuning curve was plotted to determine the cell’s directional tuning. HDs were sorted into one of sixty 6° bins. On the basis of the cell’s firing rate vs. HD tuning curve, we monitored the cell’s 1) preferred firing direction (the HD at which the cell fired maximally), 2) peak firing rate (the bin with the maximal firing rate), 3) background firing rate (the mean firing rate from all bins that were ±18° away from the cell’s directional firing range), 4) directional firing range (the range of head directions in which the firing rate was elevated above background level (based on a triangular model, Taube et al. 1990a), and 5) directional information content (IC), which was calculated as:

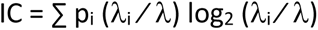

where p_i_ is the time spent with the head pointing in the ith bin divided by the total time (probability that the head pointed in the ith bin); λ is the mean firing rate of the cell in the ith bin; and λ_i_ is the overall firing rate of the cell for the entire recording session. We also computed the mean vector *R* using circular statistics; this value varies between 0 and 1, with 0 indicating no directional variation and 1 indicating perfect directional variation (Batschelet 1981). To determine whether cell firing was significantly modulated by the animal’s directional heading, we performed both a Rayleigh test based on the computed mean vector length *R* (Batschelet, 1981) and a shuffle test. For the Rayleigh test, the critical significance level of *R* is determined by the number of observations (which was defined as the sum of all firing rates from the 60 directional bins), and if the *R* value meets this significance level the distribution is considered to be non-random. Cells that exhibited significantly non-random distributions were considered directionally tuned. HD cells typically have Rayleigh r values ≥ 0.5, although many cells in this study had lower *R* values because the cells appeared more HD-modulated rather than ‘classical’ HD cells.

To aid in determining whether a cell was directionally modulated we also performed a shuffle test on its HD vs. For the shuffle test we shuffled the time series of spikes relative to the HD series and recomputed the mean vector length *R.* We shuffled the time series 500 times and computed the *R* value for each shift. We then ranked all the *R* values and if the *R* value from the non-shifted time series fell above the 99^th^ percentile, the cell was considered directionally modulated.

We also constructed HD tuning curves based on whether the animal was turning CW or CCW. These tuning curves were constructed using the same method as described above, but was confined to samples based on either the CW or CCW turn directions and when the animal was turning its head ≥ 45°/sec.

### Angular head velocity

Angular head velocity (AHV) was calculated by taking the first derivative of the HD time series, which was first smoothed by calculating a five-point running average. After smoothing, instantaneous AHV was calculated for each time point as the slope of the best-fit least-squares line fitted to the current time point, the two preceding time points, and the two subsequent time points. The slope of this best-fit line was defined as the AHV for the center point sample of the five sample window. Positive angular velocities corresponded to CCW head turns while negative angular velocities corresponded to CW head turns. The resulting AHV time series was then matched with the spike time series to construct AHV tuning plots (firing rate vs. AHV). Samples were sorted into 6°/sec bins within the range of 0-204°/sec for both CW and CCW head turns. Only bins with at least 30 samples (0.5 sec) were included in the plot.

AHV tuning curve parameters: From the firing rate/AHV functions, the following parameters were calculated:

1) Baseline firing rate: defined as the mean firing rate when the rat’s AHV was near zero (the mean value averaged across the 0 to ±6°/sec bins);
2) Correlation (Pearson’s *r*): the correlation coefficient for the best-fit lines of the CW and CCW portions of the AHV tuning curve; CCW turns were assigned positive values and CW values assigned negative values;
3) Slope: the slope of the best-fit line for the CW and CCW portions of the tuning curve, using absolute values for the CW slopes.

The AHV tuning curve was divided into four ranges based on AHV: (CW, 0–90°/sec and CW, 90– 204°/sec; CCW, 0–90°/sec and CCW, 90–204°/sec). A best-fit line and corresponding Pearson’s *r* value were calculated for each range. Because most head turns occurred at velocities < 90°/sec, this portion of the AHV tuning curve provided the most reliable sampling. Therefore, both 0-90°/s ranges and the 0–204°/s ranges, were used for classifying cells as AHV sensitive (see below).

### Cell classification for AHV

Two methods were used for classifying cells as sensitive to AHV: 1) meeting a certain threshold for correlation and slope values from the firing rate vs. AHV plot; 2) passing a shuffled test for these same criteria. For each cell we plotted two types of firing rate vs. AHV tuning curves:

1) HD-independent: Tuning curves that included all AHV samples regardless of the animal’s HD,
2) HD-dependent: Tuning curves that used AHV samples restricted to within ±60° of the cell’s preferred firing direction.

Cells were classified as tuned to AHV if they met the following three criteria for either the HD-independent or HD-dependent AHV tuning curves:

1) Pearson’s *r* correlation ≥ 0.5 for either the CW or CCW portion of the tuning curve,
2) Slope ≥ 0.05 spikes/deg for this same best-fit line,
3) Correlation and slope values for the best-fit line passed a 500 sample shuffle test, where both values had to exceed the 99^th^ percentile of the shuffled distribution.

Cells were classified as AHV-sensitive if they met these criteria for either the 0-90°/sec or 0-204°/sec ranges. Further, the cell had to pass this set of criteria for both the first and second halves of the recording session to ensure stability.

### AHV Score

To measure the strength of AHV sensitivity we devised an AHV score for each cell by fitting a best-fit line through each of the CW and CCW portions of the AHV tuning curve. We then multiplied the correlation value of these best-fit lines by its corresponding slope. The absolute value for each measure (CW and CCW) was compared with one another and the maximum value was multiplied by 100. This procedure was conducted for both the 0-90 and 0-204°/sec ranges and the highest value across these two ranges was defined as the cell’s AHV score. AHV scores for all HD cells ranged from 0 to 30, with AHV-dependent cells having scores ≥ 3.0.

### Linear velocity

Linear velocity was calculated by first computing the instantaneous speed of the animal for each 1/60th sec sample. This method was accomplished by fitting a best-fit line over a five-sample window for both the x and y dimensions (Bassett et al., 2007). The slopes of these lines were represented by the changes in the x and y dimensions, respectively, and the instantaneous speed at the center time point of the window was defined as the square root of x^2^ + y^2^. We then constructed a LV tuning curve by sorting all samples into 1 cm/sec bins and applied an AHV filter to exclude all samples where the rat was turning its head > 10 °/sec. This approach enabled us to filter out samples that could be a confound to analyzing samples for LV. The spikes associated with each sample were similarly sorted, summed for each LV bin, and divided by the total time in that bin to yield the average firing rate. A firing rate versus LV plot was then created, and a best-fit linear line was computed for the range of 0 - 30 cm/s.

From this best-fit line we calculated the correlation value (Pearson’s *r*) and slope. An LV score was computed for each cell by multiplying the absolute value of the correlation and slope values by 100 (similar to the AHV score calculation described above). For a cell to be classified as sensitive to LV it had to meet the following criteria:

1) LV score ≥ 10,
2) Correlation value (Pearson’s *r*) of the best-fit line for its tuning curve ≥ 0.7 and slope ≥ 0.1 spike/cm,
3) The correlation value had to pass a 500 sample shuffle test at the 95^th^ percentile,
4) The correlation value between the first and second halves of the session exceeded 0.5.

We note that defining the LV score in this manner inherently favors cells with high peak firing rates due to slope values, but normalizing values would exclude many cells that were clearly sensitive to LV.

In performing the LV analyses, we found that a large proportion of AHV-dependent cells were also classified as LV-sensitive. This finding raised concern that this result might be due to an artifact of tracking the rat’s movement via a two-spot tracker. We therefore performed an additional analytical method, where the rat’s location was averaged over 250 msec intervals instead of 1/60^th^ s samples, and thereby reducing the impact of small positional changes. This method yielded results that were consistent with the original method (data not shown), suggesting that the large proportion of cells that were also classified as LV sensitive is unlikely to be due to an artifact of the tracking approach.

### Statistics

All mean values reported include the SEM. One-way ANOVA tests were used to compare: 1) HD cell properties between AHV-dependent and AHV-independent cells within a brain region (LMN and DTN), and 2) properties across different brain areas (LMN, DTN). For each analysis, a Tukey’s HSD post-hoc test was conducted to explore significant main effects. A Chi-square test was used to test whether the proportion of AHV or LV cell type differed significantly across brain regions – LMN and DTN. A Yates correction was not applied because the sample sizes were sufficiently large to make the correction unnecessary. All tests used an α level of 0.05.

### Histology

Upon completion of the experiments, all animals were anesthetized deeply with sodium pentobarbital (100 mg/ml, i.p.) and a small anodal current (20 µA, 10 s) was passed through one of the electrodes to later conduct a Prussian blue reaction in the tissue. The animals were then perfused transcardially with 0.9% saline (2 min) followed by 10% formalin in 0,9% saline (15 min). The brains were placed in 10% formalin for at least 48 h. Two days prior to brain sectioning, potassium ferrocyanide was added to the 10% formalin mix for 24 h, and then the brain was placed in 20% sucrose for another 24 h. The brains were frozen and sectioned (30 µm) in the coronal plane. The sections were placed on gelatin-coated microscopic slides and after allowing 1-2 days for drying were stained with thionin and then cover slipped and allowed to dry again. All brains were then examined through a light microscope to determine the location of the marking lesion and electrode tracks. Histological analyses confirmed that 27 animals had electrodes that passed through the LMN and 10 animals had electrodes that were localized to the DTN.

## Acknowledgements

We would like to thank all the Undergraduate students who, over the years, have recorded many of the anterodorsal thalamic HD cells analyzed in this paper. We also thank Matthijs van der Meer for helpful discussions on how HD cell attractor networks work and Juliana Taube for help with constructing some of the figures. This work was supported by National Institutes of Health (USA) - National Institute of Neurological Disorders and Stroke Grants NS053907 and NS111695.

## Supplementary Figure Legends

**Supplementary Figure 1.** HD-independent vs. HD-dependent AHV graphs. Each row is one cell. The first column shows the HD x firing rate tuning curve. The second column depicts the HD-independent AHV tuning curve. The third column depicts the HD-dependent AHV tuning curve. **A, B)** AHV tuning curves were usually similar between HD-independent and HD-dependent graphs. **C, D)** Two examples where the AHV tuning curves were not similar between the two graphs. In *C*, the cell went from a Symmetric cell for the HD-independent tuning curve to a mildly AHV-sensitive Asymmetric-unresponsive cell for its entire tuning curve. In *D*, the cell was classified as an Asymmetric-unresponsive cell based on its HD-independent AHV tuning curve, but was only mildly sensitive to AHV based on its HD-dependent AHV tuning curve. For all AHV graphs, note that the HD-dependent graph has a higher firing rate than the HD-independent graph. All axes are labelled as shown in the top row.

**Supplementary Figure 2.** Relationship between the direction of AHV sensitivity (CW vs. CCW) and the hemisphere where the cells were recorded. **A-C)** Three LMN HD+AHV Asymmetric cells that were all recorded in the same animal from the right hemisphere. Note that all three cells were sensitive to CCW head turns, which is opposite to that observed from all the Asymmetric cells from other animals where AHV sensitivity was in the direction towards the hemisphere ipsilateral to the implant; see examples in Figure 3. Each row is one cell. Formatted columns are the same as Figure 2. All axes are labelled as shown in the top row.

