## Supplemental Figure 1 for "Generating the head direction signal: Two types of head direction cells in the lateral mammillary nuclei and dorsal tegmental nuclei"

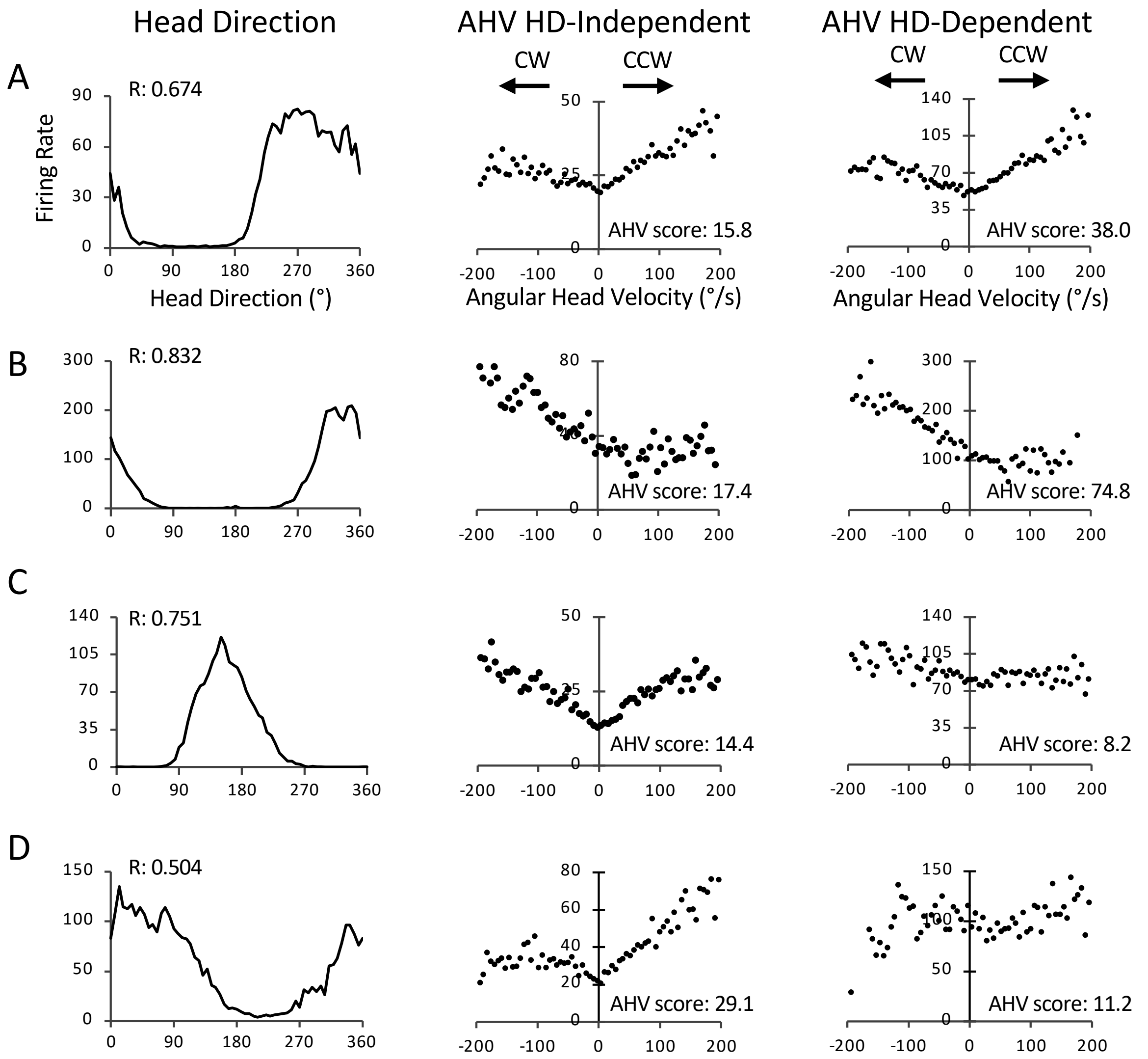


**Supplementary Figure 1**. HD-independent vs. HD-dependent AHV graphs. Each row is one cell. The first column shows the HD x firing rate tuning curve. The second column depicts the HD-independent AHV tuning curve. The third column depicts the HD-dependent AHV tuning curve. **A, B)** AHV tuning curves were usually similar between HD-independent and HD-dependent graphs. **C, D)** Two examples where the AHV tuning curves were not similar between the two graphs. In *C*, the cell went from a Symmetric cell for the HD-independent tuning curve to a mildly AHV-sensitive Asymmetric-unresponsive cell for its entire tuning curve. In *D*, the cell was classified as an Asymmetric-unresponsive cell based on its HD-independent AHV tuning curve, but was only mildly sensitive to AHV based on its HD-dependent AHV tuning curve. For all AHV graphs, note that the HD-dependent graph has a higher firing rate than the HD-independent graph. All axes are labelled as shown in the top row.

**Suppl Fig. 1**
