## Supplemental figure 2 for "Generating the head direction signal: Two types of head direction cells in the lateral mammillary nuclei and dorsal tegmental nuclei"

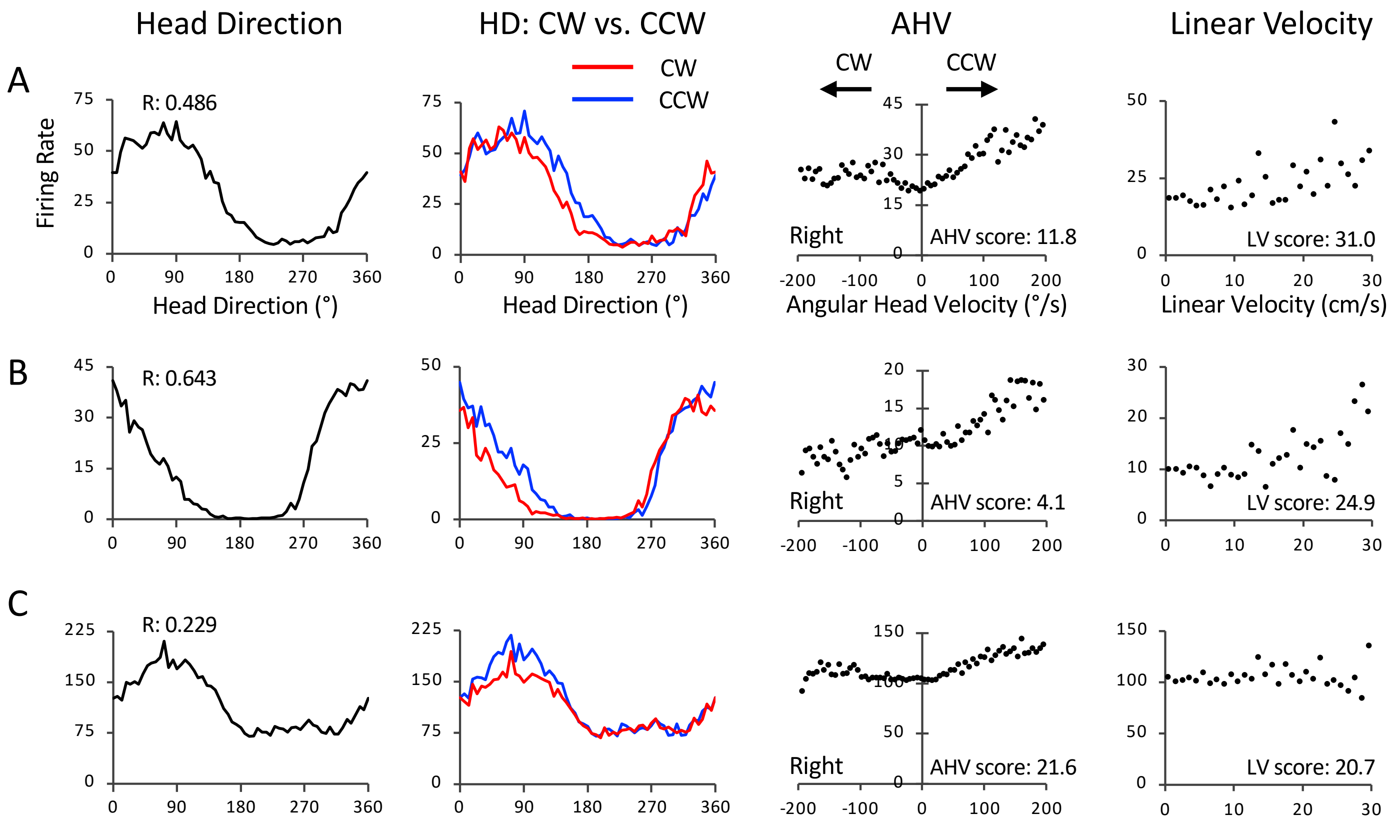


**Supplementary Figure 2**. Relationship between the direction of AHV sensitivity (CW vs. CCW) and the hemisphere where the cells were recorded. **A-C)** Three LMN HD+AHV Asymmetric cells that were all recorded in the same animal from the right hemisphere. Note that all three cells were sensitive to CCW head turns, which is opposite to that observed from all the Asymmetric cells from other animals where AHV sensitivity was in the direction towards the hemisphere ipsilateral to the implant; see examples in Figure 3. Each row is one cell. Formatted columns are the same as Figure 2. All axes are labelled as shown in the top row.

**Suppl Fig. 2**
